# Dichotic listening deficits in amblyaudia are characterized by aberrant neural oscillations in auditory cortex

**DOI:** 10.1101/2020.11.27.401604

**Authors:** Sara Momtaz, Deborah W. Moncrieff, Gavin M. Bidelman

**Affiliations:** School of Communication Sciences & Disorders, University of Memphis, Memphis, TN, USA; Institute for Intelligent Systems, University of Memphis, Memphis, TN, USA; University of Tennessee Health Sciences Center, Department of Anatomy and Neurobiology, Memphis, TN, USA

**Keywords:** Auditory processing disorders (APD), event-related brain potentials (ERPs), gamma/beta band response, hemispheric asymmetries, phase-locking, time-frequency analysis

## Abstract

Children diagnosed with auditory processing disorder (APD) show deficits in processing complex sounds that are associated with difficulties in higher-order language, learning, cognitive, and communicative functions. Amblyaudia (AMB) is a subcategory of APD characterized by abnormally large ear asymmetries in dichotic listening tasks. Here, we examined frequency-specific neural oscillations and functional connectivity via high-density EEG in children with and without AMB during passive listening of nonspeech stimuli. Time-frequency maps of these “brain rhythms” revealed stronger phase-locked beta-gamma (∼35 Hz) oscillations in AMB participants within bilateral auditory cortex for sounds presented to the right ear, suggesting a hypersynchronization and imbalance of auditory neural activity. Brain-behavior correlations revealed neural asymmetries in cortical responses predicted the larger than normal right-ear advantage seen in participants with AMB. Additionally, we found weaker functional connectivity in the AMB group from right to left auditory cortex, despite their stronger neural responses overall. Our results reveal abnormally large auditory sensory encoding and an imbalance in communication between cerebral hemispheres (ipsi-to -contralateral signaling) in AMB. These neurophysiological changes might lead to the functionally poorer behavioral capacity to integrate information between the two ears in children with AMB.

## 1. INTRODUCTION

Auditory processing disorder (APD) refers to the inability to appropriately process complex sound information that may lead to higher-order deficits such as language, learning, cognitive, and communication impairment despite normal hearing. However, there is currently no gold standard test for a differential diagnosis of APD (AAA, 2010; ASHA, 2005; BSA, 2011). The neurophysiological underpinning of APD likely represents a spectrum of varying etiologies (e.g., corpus callosum lesions, low birth weight, prematurity, chronic ear infection) (Jerger et al., 2002; Milner et al., 2018; Moncrieff, 2006) and there is limited consensus in terms of prevalence, test battery, diagnostic criteria, or even precise definition of APD (Dillon et al., 2012; Moore and Hunter, 2013; Wilson and Arnott, 2013). Moreover, common manifestations of auditory symptoms in various disorders [e.g., attention-deficit/hyperactivity disorder (ADHD), dyslexia, or specific language impairment (SLI)] may arise from the same basic auditory deficits that make the differential diagnosis of these patients even more complicated (Gilley et al., 2016; Milner et al., 2018; Riccio et al., 1994; Sharma et al., 2009). APD may also produce functional and anatomical abnormalities that are not limited solely to the auditory pathways (Farah et al., 2014; Kim et al., 2009; Micallef, 2015; Owen et al., 2013; Pluta et al., 2014; Schmithorst et al., 2011). These limitations have motivated the use of objective approaches to define specific neural mechanisms that contribute to auditory-based learning disorders including APD.

The auditory system’s neuroanatomy is bilateral and functional processing occurs primarily through crossed (i.e., contralateral) pathways. However, ipsilateral and contralateral pathways are more symmetrical compared to other modalities (Rauschecker and Scott, 2009). Amblyaudia (AMB) is a subtype of APD that emerges when listeners fail to normally process competing stimuli presented simultaneously to both ears (Moncrieff et al., 2016). AMB is characterized by a larger than normal asymmetry on two or more dichotic listening tests despite normal audiometry (Moncrieff et al., 2016), with performance in the non-dominant ear often >2 SDs below normal while performance in the dominant ear is normal. It is presumed that the listener’s dominant ear offsets and counterbalances deficiencies in binaural processing of auditory information while the non-dominant ear is developmentally deprived, leading to a deficit in binaural integration of complex signals (Lamminen and Houlihan, 2015). Children with AMB have a range of difficulties in the domains of auditory (e.g., poor verbal working memory, speech comprehension (especially in noise), localization], cognitive [e.g., attention), linguistic (e.g., syntactic impairment), and social processing (e.g., poor adaptive and self-esteem skills) (Lamminen and Houlihan, 2015; Moncrieff et al., 2004; Popescu and Polley, 2010; Whitton and Polley, 2011). A unilateral deficit during dichotic listening tasks has long been attributed to callosal dysfunction (Musiek, 1983; Musiek and Weihing, 2011), though later studies have recognized that *functional* asymmetries along the auditory system might provide a basis for the disorder, possibly as low as the brainstem superior olivary complex (Hiscock and Kinsbourne, 2011; Moncrieff et al., 2008; Tollin, 2003). Otitis media, closed-head injuries, and co-morbid disabilities in early childhood may contribute to periods of auditory deprivation that are well known to produce structural and functional abnormalities in the brainstem (Clopton and Silverman, 1977; Coleman and O’Connor, 1979; Moore and Irvine, 1981; Popescu and Polley, 2010; Silverman and Clopton, 1977; Smith et al., 1983; Webster and Webster, 1979) that may lead to AMB. With altered auditory input during development, the brain may recalibrate central auditory circuits to compensate for transient periods of hearing loss from the affected ear(s). Long-term consequences of such perceptual deficits may lead to linguistic, cognitive, and social concomitants that further outweigh this compensatory mechanism (Keating and King, 2013; Popescu and Polley, 2010), leading to poorer binaural hearing skills.

Neuroimaging studies have shed light on the physiological basis of APD and its related AMB variant. In a quasi-dichotic ERP study using speech stimuli, children with AMB were unable to process aberrant stimuli presented to their left ears with competing signals presented to their right ears (Jerger et al., 2002; Moncrieff et al., 2004). Similarly, early fMRI studies noted hemispheric deficiencies when children with AMB listened to dichotic stimuli compared to normal controls (Hashimoto et al., 2000; Moncrieff et al., 2008). Because dichotic listening places greater demands on attention, engaging more cognitive control to process information in the non-dominant ear, it typically results in greater activity in the left superior temporal gyrus, inferior frontal gyrus, and superior anterior cingulate (Hugdahl et al., 2003; Jäncke et al., 2001; Jäncke and Steinmetz, 2003). Compared to the high levels of neural activation prior to therapy, activations in children with AMB were significantly reduced following auditory therapy designed to enhance non-dominant ear performance, especially in bilateral auditory cortices, precentral gyrus and in right hemisphere regions known to be involved in semantic processing and working memory (Moncrieff & Schmithorst, in review). Despite the appropriate spatial resolution of fMRI, the technique is severely limited in time resolution (∼2-3 s) that is required for precise auditory temporal processing (Cantiani et al., 2019; Gaudet et al., 2020). While ERPs offer fine timing precision, volume conduction and referential recordings cannot uncover the brain mechanisms contributing to changes in scalp-recorded neural activity (Cantiani et al., 2019). Moreover, conventional EEG approaches neglect potentially important rhythmic, spectrotemporal details of brain activity that could provide further insight into the hearing function and its disorders (Makeig, 1993; Nunez and Srinivasan, 2006). In this vein, neural oscillations have elucidated new perspectives on the underlying functional networks of the auditory processing hierarchy that mediate complex behaviors (Gilley et al., 2016).

Phase-locked (evoked) and non-phase locked (induced) responses are two major components of EEG activity (brain oscillations) that represent different aspects of perception-cognition and the auditory processing hierarchy (Bidelman, 2015; Giraud and Poeppel, 2012; Shahin et al., 2009). Neural oscillations are carried within different frequency bands of the EEG [delta (1–3 Hz), theta (4–8 Hz), alpha (9-12), beta (13–30 Hz), low gamma (30–70 Hz), and high gamma (70–150 Hz)], though they may be lower in young children (Saby and Marshall, 2012). These brain rhythms provide a unique insight into auditory perceptual processing not available in conventional ERP measures (Bidelman, 2015). For instance, β band reflects rhythmic processing (Gilley et al., 2016), sensory integration (Brovelli et al., 2004), working memory (Shahin et al., 2009; Zarahn et al., 2007), auditory template matching (Bidelman, 2015, 2017; Shahin et al., 2009; Yellamsetty and Bidelman, 2018), and novelty detection (Cope et al., 2017; HajiHosseini et al., 2012; Sedley et al., 2016). γ band is associated with early feature selection (Gilley et al., 2016), local network synchronization (Giraud and Poeppel, 2012), auditory object selection (Tallon-Baudry and Bertrand, 1999), attention (Gilley et al., 2016), as well as top-down and bottom-up auditory integration (Tallon-Baudry and Bertrand, 1999; Trainor et al., 2009).

Here, we exploited the rich spectrotemporal details of EEG to investigate changes in neural oscillatory activity in children with AMB. To better understand the underlying physiology of auditory processing deficits related to AMB, we evaluated multichannel EEGs in typically developing children and age-matched peers diagnosed with AMB using passively evoked, non-speech stimuli. With source and functional connectivity analysis, we analyzed the time-frequency information of neural oscillations stemming from the auditory cortex (AC). Fine-grained source analysis enabled us to focus on the underlying neural substrates that account for changes in scalp-level data (Gaudet et al., 2020). We hypothesized brain rhythms might differ in children with dichotic listening problems. Based on previous work (Gilley et al., 2016), we predicted increased β or γ band auditory cortical activity might convey an imbalance in neural representation predictive of the stark ear asymmetries observed in AMB children at the behavioral level. We further posited that AMBs might show a functional imbalance in communication (connectivity) between cerebral hemispheres consistent with the central transmission hypothesis of AMB.

## 2. MATERIALS & METHODS

### 2.1 Participants

Twenty-six children ranging in age from 9-12 years were divided into two groups based on their behavioral scores on dichotic listening (DL) tests used as part of a comprehensive assessment of auditory processing (described below). Children with an asymmetric pattern of results from the DL tests were placed into the AMB group (n=14); the remaining children were within normal limits (WNL) (n=12). Groups were matched in age (AMB: 10.14 ± 1.67 years, WNL: 10.75 ± 1.05 years, *t*_24_ = -1.38, p = 0.18) and gender (AMB: 10/4 male/female; WNL 8/4 male/female; Fisher exact test, p = 1). All participants had normal health at the time of study without any history of neurological impairment, head injury, chronic disease, or hearing loss (≤25 dB HL screened from 500-4000 Hz; octave frequencies). They were recruited either from APD evaluation clinic referrals or by fliers distributed throughout the community. Parents of participants gave written informed consent in compliance with a protocol approved by the Institutional Review Board at the University of Pittsburgh.

### 2.2 Behavioral evaluation

Three DL tests were performed at 50 dB HL to measure participants’ binaural listening skills and laterality: Randomized Dichotic Digit Test (RDDT) (Moncrieff and Wilson, 2009), Dichotic Words Test (DWT) (Moncrieff, 2015), and Competing Words subtest from the SCAN-C (CW) (Keith, 1986).

#### RDDT

The Randomized Dichotic Digit Test (RDDT) was performed with single syllable digits from 1-10 (except 7) that were recorded with a male voice (Moncrieff and Wilson, 2009). Children were instructed to repeat both digits presented dichotically to the right and left ears (order did not matter). They were instructed to guess if they were unsure of what they heard. There were 18 presentations each of single pairs (n=18 digits), double pairs (n=36 digits), and triple pairs (n=54 digits). Correct percent recall was scored for each ear separately and interaural asymmetry was calculated as the difference in performance between the two ears. The scores for the double pair condition were compared to normative data for the test.

#### DWT

The DWT presents duration-matched single-syllable words dichotically via headphones (Moncrieff, 2015). Like the RDDT, the patient was asked to repeat both words. Correct percent recall was scored for each ear separately and interaural asymmetry was calculated as the difference in performance between the two ears.

#### CW

Competing words (CW) is a subtest of the SCAN, Screening Test for Auditory Processing Disorder (Keith, 1986). The test includes 30 pairs of single-syllable words recorded by a natural male voice. Stimuli were presented with the same onset time in both ears. Correct percent word recall was scored for each ear separately during two separate conditions in which the listener was asked to repeat the right ear first or the left ear first.

Individual ear scores were converted from the right and left to dominant and non-dominant so that interaural asymmetry was always a positive number reflective of the difference in performance between the two ears. AMB is characterized by an abnormally large interaural asymmetry and is diagnosed when a larger than normal interaural asymmetry is observed on at least two dichotic listening tests (Moncrieff et al., 2016).

### 2.3 EEG recording procedure

#### Stimuli

Auditory electrophysiological responses were elicited using a 385 μsec biphasic acoustic click (first phase = 181 μs, second phase = 204 μs). Stimuli were presented monaurally to each ear (separate runs) at 70 dB nHL via ER-3A insert earphones. The rate of the presentation was 8.5/sec (i.e., 117 ms interstimulus interval) and 1000 sweeps were collected for each ear of presentation.

#### EEG

Participants were seated in an electrically shielded, sound-attenuating booth during EEG testing. They were asked to close or relax their eyes (e.g., focus on something in their lap). EEGs were recorded using a 32-channel electrode cap (Neuroscan QuikCap) that was held in place using a chin strap. Electrode positions in the array followed the international 10-20 system (Oostenveld and Praamstra, 2001). Each Ag/AgCl electrode was filled with a water-soluble conductive gel to achieve impedances <5 kΩ. EEGs were recorded using the Neuroscan Synamp^2^ at a sample rate of 10 kHz. During recording, electrodes were referenced to linked earlobes with the ground electrode placed at the midforehead (AFz). Data were re-referenced to the common average offline for subsequent analysis.

We used BESA Research 7.0 (BESA, GmbH) to preprocess the continuous EEG data. Recordings were epoched into single trials from -10 ms to 100 ms, bandpass filtered from 10 to 2000 Hz, and baseline corrected to the pre-stimulus interval. These parameters were designed to capture middle latency responses (MLR) of the auditory evoked potentials (to be reported elsewhere).

### 2.4 EEG time-frequency analysis

Prior to time-frequency analysis (TFA), we further cleaned the EEG data of artifactual segments (e.g., blinks). Paroxysmal electrodes were spherically spline interpolated. We then used a two-pronged approach for artifact rejection. Trials exceeding ±500 µV were first rejected using thresholding. This was followed by a gradient criterion, which discarded epochs containing amplitude jumps of > 75 µV between any two consecutive samples. This resulted in between 860 – 1000 artifact-free trials for analysis. It should be noted this number of sweeps is much greater (∼10x) than what is typically required for TFA (Shahin et al., 2010; Yellamsetty and Bidelman, 2018). Critically, trial counts did not differ between groups for either left (*t*_25_=0.19, *p*=0.85) or right (*t*_25_=0.19, *p*=0.74) ear recordings indicating similar overall signal-to-noise ratio.

We transformed each listeners’ scalp potentials into source space using BESA’s Auditory Evoked Potential (AEP) virtual source montage (Bidelman, 2017; Mankel et al., 2020; Scherg et al., 2002). This applied a spatial filter to all electrodes that calculated their weighted contribution to the scalp recordings. The AEP model includes 11 regional dipoles distributed across the brain including bilateral AC [Talairach coordinates (*x,y,z*; in mm): *left* = (−37, -18, 17) and *right* = (37, -18, 17)]. Regional sources consist of three dipoles describing current flow (units nAm) in the radial, tangential, and anterior-posterior planes. We extracted the time courses of the radial and tangential components for left and right AC sources as these orientations capture the majority of variance describing the auditory cortical ERPs (Picton et al., 1999). Orientations were pooled in subsequent analyses. This approach allowed us to reduce each listeners’ 32 channel data to 2 source channels describing neuronal activity localized to the left and right AC (Mankel et al., 2020; Price et al., 2019).

We then performed a TFA on the source data to evaluate frequency-specific differences in neural oscillations between groups (Hoechstetter et al., 2004)^1^. From single-trial epochs, we computed a time-frequency transformation using a sliding window analysis (complex demodulation; Papp and Ktonas, 1977) and 20 ms/2.5 Hz resolution step sizes. These settings permitted analysis of frequencies 10-80 Hz across the entire epoch window. The resulting spectral maps were then produced by computing inter-trial phase-locking (ITPL) at each time-frequency point across single trials (Hoechstetter et al., 2004). These maps are three-dimensional functions (time x frequency x ITPL), akin to neural spectrograms (see Fig. 2) that visualize ITPL (phase-locking strength) rather than raw amplitudes. ITPL varies between 0-1 in which 0 reflects stochastic noise (i.e., absence of repeatable brain activity) and 1 reflects perfect trial-to-trial response repeatability (Tallon-Baudry et al., 1996). ITPL maps reflect the change in neural synchronization relative to baseline (−10 to 0 ms) and contain evoked neural activity that is time- and phase-locked to the eliciting repetitive click stimulus (Bidelman, 2015; Shahin et al., 2010). Maps were upsampled by a factor of 10x (bicubic interpolated) in the time and frequency dimensions for further visualization and quantification.

**Figure 1:**
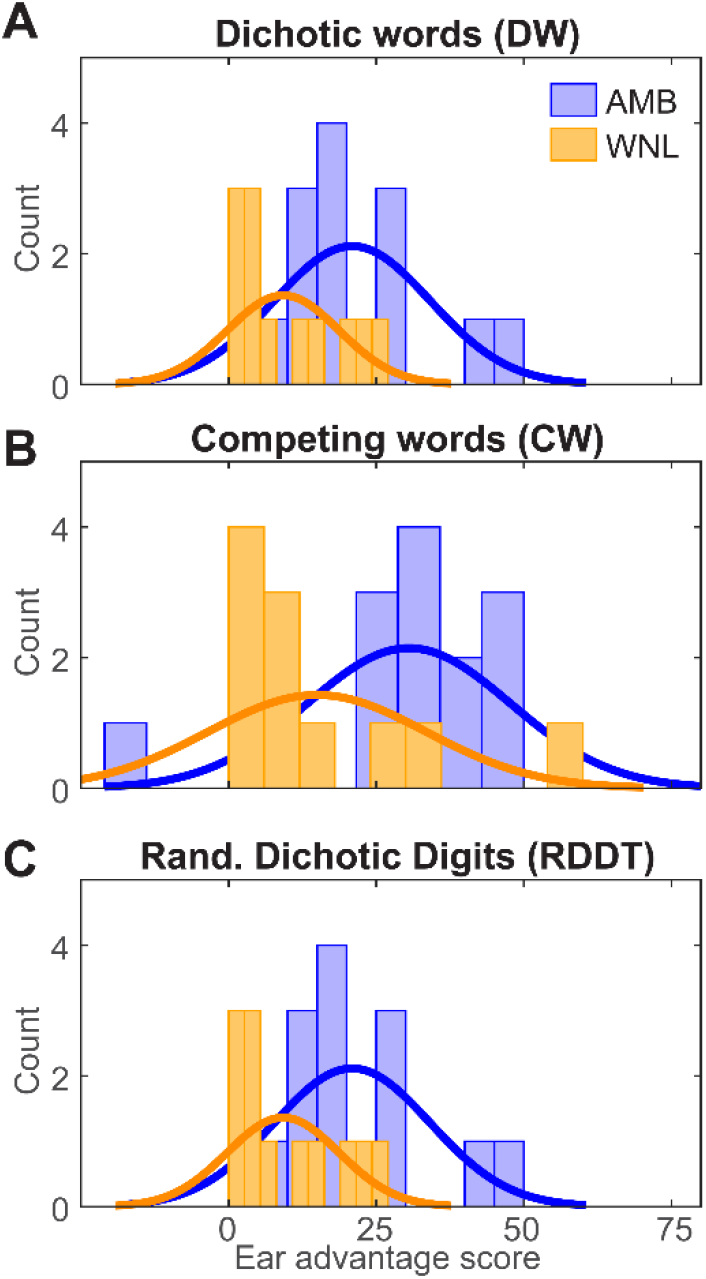
Group histograms of behavioral ear advantage scores in dichotic listening. (**A**) Dichotic words (DW) test. (**B**) Competing words (CW) test. (**C**) Random Dichotic Digits Test (RDDT). AMBs show stronger ear advantage scores (i.e., larger asymmetry) for all three tasks. Solid lines denote normal curve fits.

**Figure 2:**
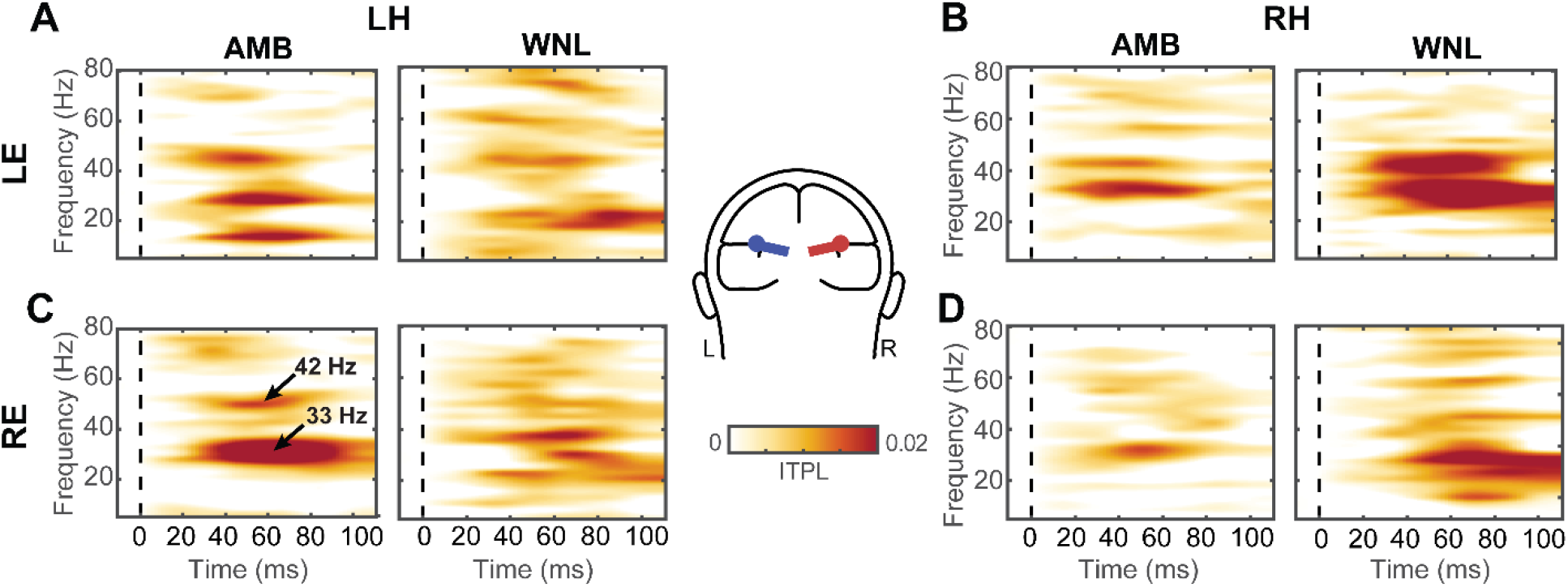
Time-frequency maps of neural oscillatory activity from auditory cortex across groups, hemispheres, and ears. (**A, C**) Left hemisphere (LH) responses. (**B, D**) Right hemisphere (RH) responses. Only maps for the radial dipole orientation are shown for clarity (inset). ITPL maps reflect “evoked” variations of phase-locked EEG relative to baseline (i.e., power spectrum of the ERP). *t*=0, click stimulus onset. Time-frequency maps reveal strong neural synchrony near the ∼40 Hz frequency band that is modulated by group membership, cerebral hemisphere, and ear of stimulus presentation. Red colors denote stronger neural phase synchrony across trials. LE/RE, left/right ear; LH/RH, left/right hemisphere.

Initial visual inspection of ITPL spectrograms revealed prominent group differences in the high-β/low-γ frequency band (i.e., ∼33 Hz; see Fig. 2C). To quantify these group effects, we extracted the time course of the 33 Hz band from each spectrogram (see Fig. 3) per hemispheric source and ear of presentation. We then measured the peak maximum amplitude and latency from each band response manually using MATLAB 2019 (The MathWorks, Inc). Peak maxima were marked twice for each waveform. Inter-rater agreement was excellent (*r*=0.96). Consequently, we averaged the two independent measurements to compare band amplitudes and latencies between AMB and WNL groups.

**Figure 3:**
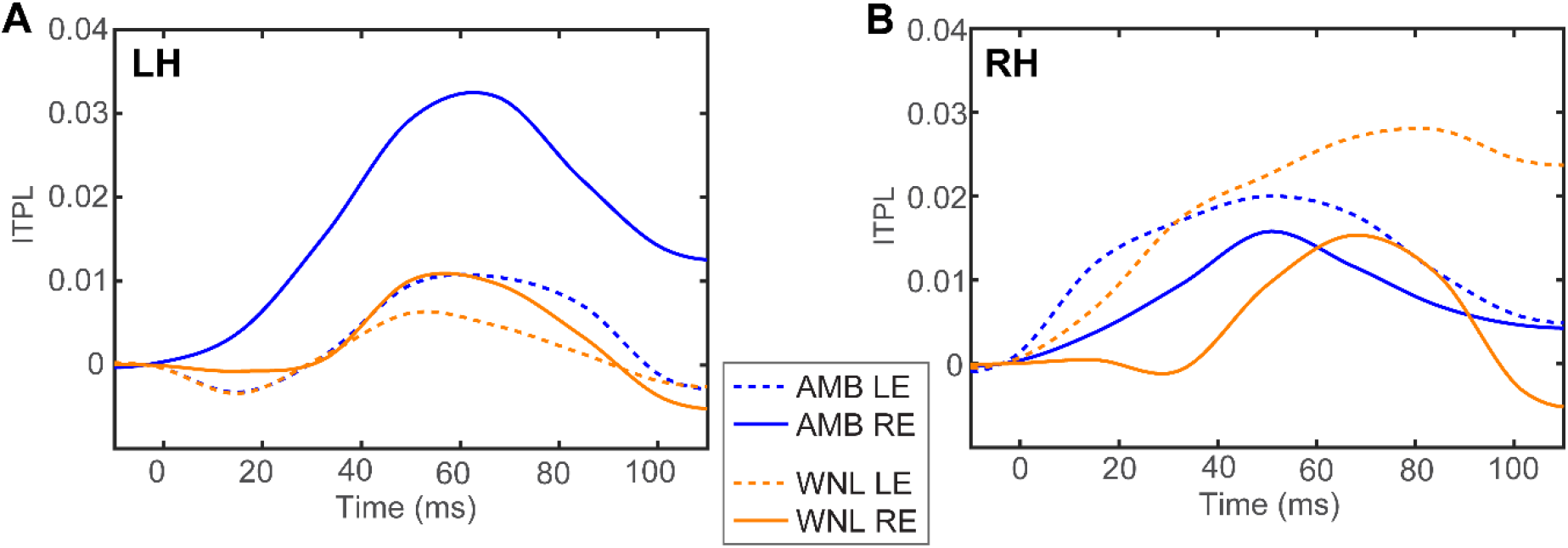
Band time waveforms from (A) left and (B) right hemisphere. Time-courses depict the temporal dynamics of phase-locking within the β/γ frequency band (i.e., 33 Hz spectral slice of Fig. 2 time-frequency maps). Note the stronger responses for right ear stimulation in AMBs.

### 2.5 Functional connectivity

We measured directed information flow between the left and right AC responses using phase transfer entropy (PTE) (Lobier et al., 2014). PTE connectivity is sensitive to hearing pathologies and individual differences in complex listening skills (Bidelman et al., 2018; Bidelman et al., 2019). It is a non-parametric measure of directed signal interaction that can be computed bi-directionally between pairs of sources (X→Y vs. Y→X) to infer causal flow between brain regions. PTE was estimated using the time series of the instantaneous phases of pairwise signals (i.e., LH and RH band waveforms) (for details, see Bidelman et al., 2019; Hillebrand et al., 2016; Lobier et al., 2014) using Otnes histogram binning (Otnes and Enochson, 1972) and a delay parameter based on the frequency content of the signal—as implemented using the *PhaseTE_MF* function in the Brainstorm toolbox (Tadel et al., 2011). This implementation assesses the statistical dependency between inter-hemispheric waveform morphology across the entire epoch window and is not time-resolved, *per se*. We computed PTE in both directions between LH and RH source β-band waveforms to quantify the relative weighting of connectivity between hemispheres per group and ear of presentation.

### 2.6 Statistical analysis

We used 2×2×2 mixed model ANOVAs (GLIMMIX, SAS ® 9.4, SAS Institute; Cary, NC) to assess all dependent variables of interest. Fixed factors were group (2 levels: WNL, AMB), ear (2 levels: LE, RE), and hemisphere (2 levels, LH, RH); subjects served as a random effect. The data were normally distributed. The significance level was set at α= 0.05.

We used correlations (Pearson’s-*r*) to evaluate relationships between neural oscillations and behavior (i.e., dichotic listening scores). For these analyses, we derived a laterality index for the neural measures, computed as the *difference* in peak ITPL between ears (i.e., laterality = ITPL_RE_ - ITPL_LE_). LH and RH responses were pooled given the lack of hemisphere effect in the omnibus ANOVAs (see *Section 3*.*2*). Neural laterality was then regressed against listeners’ ear advantage score (per RDDT, DW, and CW test), computed as the magnitude differences in behavioral performance between their dominant and non-dominant ear. This ear advantage score reflects the fact that AMB is characterized by larger than normal asymmetries in dichotic listening between ears (Moncrieff et al., 2016; Moncrieff et al., 2008).

## 3. RESULTS

### 3.1 Behavioral data

**Figure 1** shows dichotic listening performance between groups. Values indicate ear advantage, measured as the difference in behavioral performance between non-dominant and dominant ears. Regardless of group, most individuals showed a right ear advantage (AMB:11/14 = 79%, WNL:10/12 = 83%). Participants with AMB showed larger ear advantage compared to WNL listeners in all three dichotic tests including DW [AMB: 29.43 ± 20.85, WNL: 11.33 ± 11.55; *t*_24_ = 2.67, *p* = 0.013], CW [AMB: 30.54 ± 17.16, WNL: 15 ± 18.43; *t*_22_ = 2.14, *p* = 0.044], and RDDT [AMB: 20.93 ± 13.19, WNL: 9.25 ± 9.48; *t*_24_ = 2.55, *p* = 0.017]. These findings confirm large interaural asymmetry in AMB compared to the WNL group (Moncrieff et al., 2016).

### 3.2 EEG time-frequency data

ITPL spectral maps across groups, hemispheres, and ears are shown in **Figure 2**. Oscillatory responses represent evoked activity that is phase-locked to the stimulus presentation, localized here to the AC. Visual inspection of spectral activity between AMB and WNL groups showed differences in the high-β/low-γ frequency bands (i.e., 33 and 42 Hz; Fig. 2C) roughly ∼40-80 ms post-stimulation. These data expose different neural oscillatory representations of rapid auditory stimuli between groups.

To quantify these group differences, we extracted band time courses within the 33 Hz band, where group, ear, and hemispheric differences were prominent in the spectrographic maps. ITPL time courses are shown in **Figure 3**. Peak amplitudes of these waveforms are shown in **Figure 4**.

**Figure 4:**
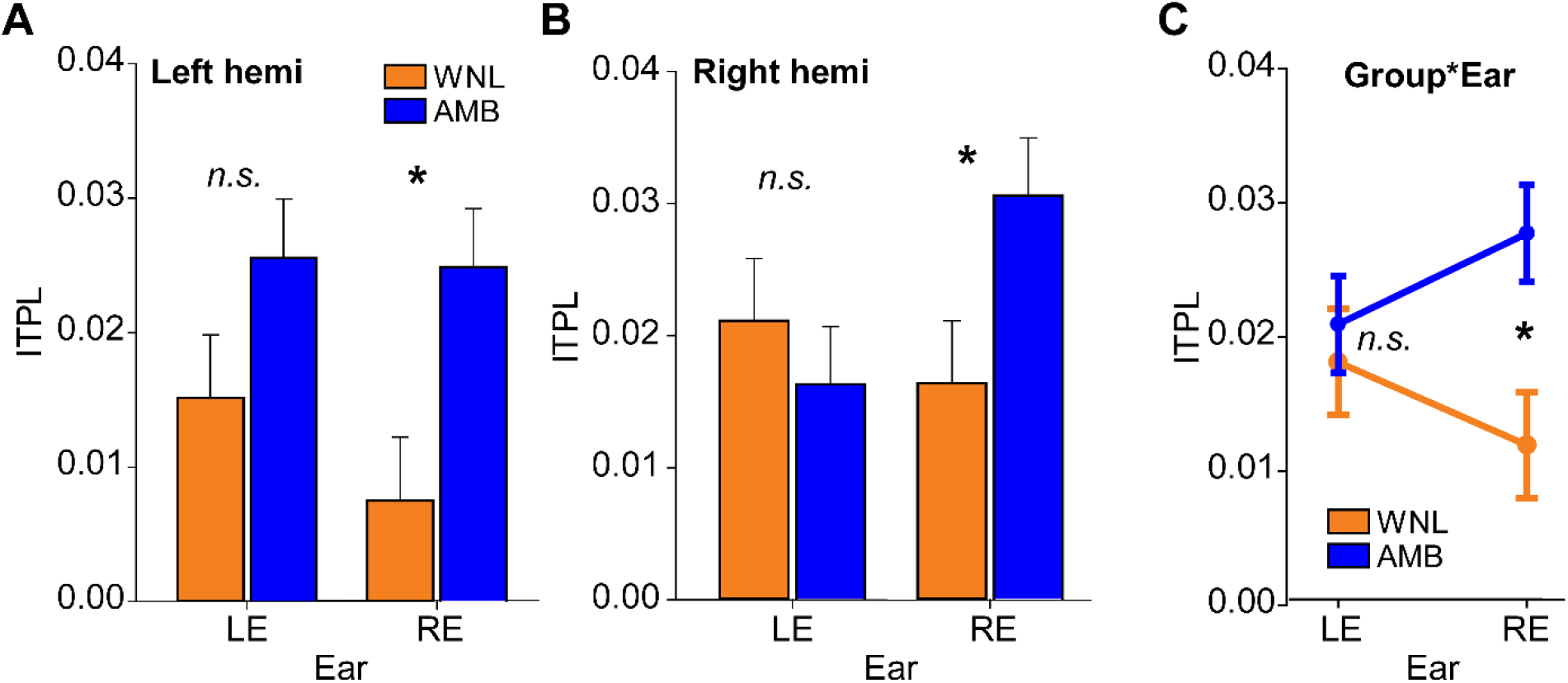
Band quantification illustrates larger neural oscillations in AMB and the right ear. (**A, B**) Peak ITPL amplitudes per group, ear, and hemisphere extracted from the time-courses shown in Fig. 3. ITPLs vary with group and ear but not hemisphere. (**C**) Group x ear interaction effect. AMBs show stronger oscillatory responses for RE stimulation. Error bars = ± 1 s.e.m., \**p*< 0.05.

An ANOVA conducted on ITPL peak amplitudes revealed a group x ear interaction in neural oscillation strength [*F*_*1,72*_ = 6.43, *p* = 0.0134]. Tukey-Kramer-adjusted multiple comparisons revealed the interaction was due to stronger amplitudes in AMB vs. WNL listeners for the right ear (*p* = 0.0041) but not the left ear (*p* = 0.60) presentation (**Figure 4C**). By group, responses were marginally larger in right vs. left ear for AMBs (*p* = 0.055) but were invariant in WNLs (*p* = 0.10). The group x hemisphere effect approached but was not significant [*F*_*1,72*_ = 3.23, *p* = 0.0764]. No other main or interaction effects with hemisphere were observed. Response latencies uniformly peaked at ∼60 ms and were invariant across groups, hemisphere, and ears (all *p*s >0.17). Thus, ∼60 ms after stimulus presentation, AMBs showed higher phase-locked high-β/low-γ responses for right ear stimulation (see Fig. 3) across both hemispheres.

### 3.3 Brain-behavior relationships between neural oscillations and dichotic listening

We evaluated the correspondence between neural ear laterality and all three behavioral ear advantage scores using correlational analysis (**Figure 5**). Correlation results revealed that the degree of ear asymmetry in neural oscillation strength (i.e., peak ITPL_RE_ - ITPL_LE_) was associated with behavioral performance on the DW test (*r* = 0.43, *p* = 0.029), whereby stronger right ear responses predicted larger ear advantage in dichotic listening. No other correlations were significant.

**Figure 5:**
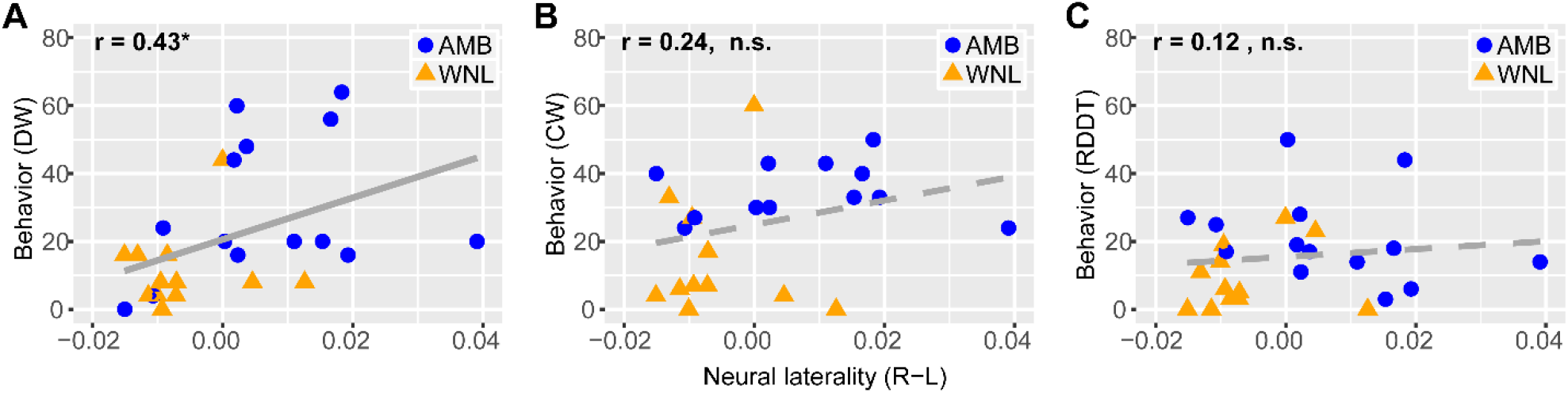
Dichotic listening deficits in AMB are associated with abnormally large asymmetries in AC oscillations. Scatters show brain-behavior correlations between ear advantage in dichotic listening (i.e., Fig. 1) and neural laterality (i.e., Fig 4; laterality = ITPL_RE_ - ITPL_LE_; pooling hemispheres). (**A**) Dichotic word (DW) test. (**B**) Competing words (CW) test. (**C**) Random Dichotic Digits Test (RDDT). Only DW scores correlate with neural responses. Stronger high-β/low-γ oscillations in the right ear (as in AMBs) predict larger ear asymmetry in DW. Solid lines = significant correlations (\**p*< 0.05), dotted lines = insignificant correlations (n.s.).

### 3.4 Functional connectivity

Interhemispheric connectivity in the leftward (RH→LH) and rightward (LH→RH) directions are shown in **Figure 6**. PTE values were well above zero (*t*-tests against PTE=0; *P*s< 0.0001) confirming significant (non-random) neural signaling in both directions. An ANOVA revealed a 3-way interaction between group x ear x-direction on connectivity strength [*F*_1,78_= 6.30, *p*=0.0141] (**Fig. 6A**). Tukey contrasts revealed this three-way effect was driven by differences in inter-hemispheric connectivity isolated to the AMB group for RE stimulation only. Specifically, when stimuli were delivered to the RE, AMBs showed weaker transmission directed from RH→LH compared to the reverse direction (p=0.0028). Connectivity strength did not differ for controls or for LE stimulation (within either group) (all *p*s > 0.13) (**Fig. 6B**). Thus, whereas LH→RH transmission was identical between groups, AMBs showed weaker transfer of information from right to left AC, despite their stronger neural responses overall (Figs. 4-5).

**Figure 6:**
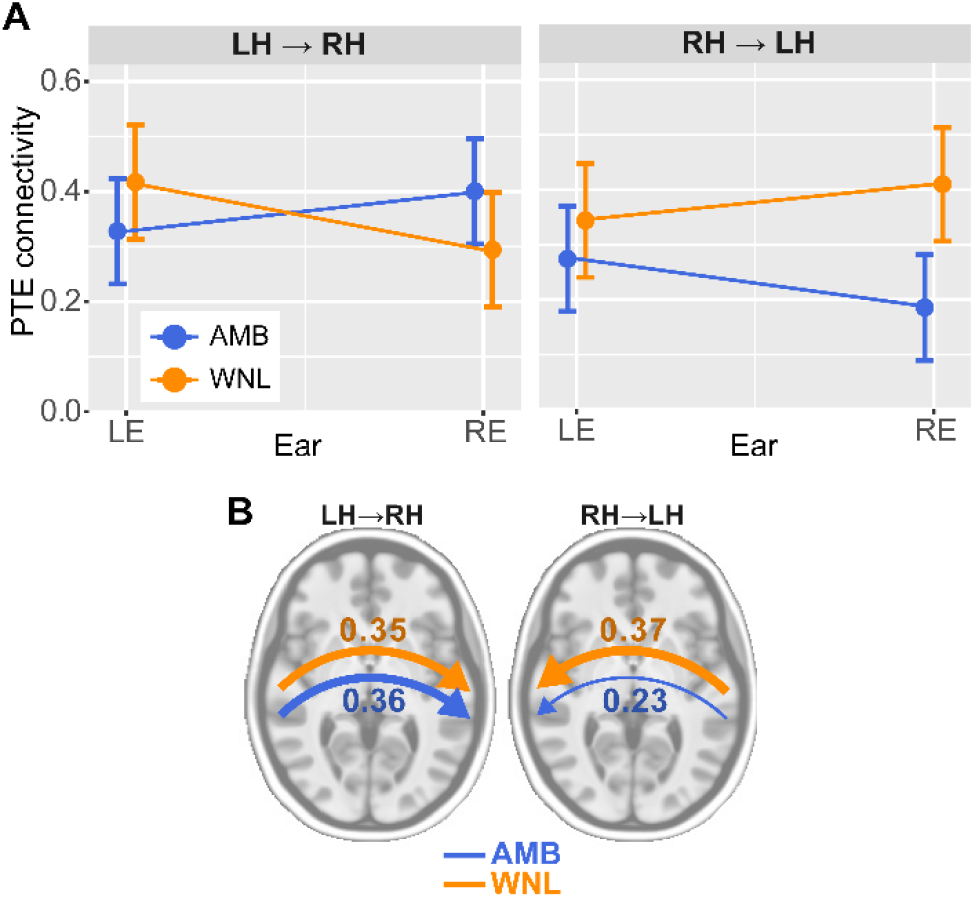
Functional connectivity directed from right to left AC is weaker in AMB listeners. (**A**) Interhemispheric connectivity measured via PTE (Lobier et al., 2014) illustrates a 3-way interaction (group x ear x-direction) on neural transmission strength. Each panel reflects interhemispheric signaling between bilateral AC directed in the left-to-right (LH→RH) or right-to-left (RH→LH) direction. (**B**) LH→RH signaling is similar between groups. Connectivity from RH→LH is weaker in AMBs for RE (but not LE) stimulation. errorbars = ±1 s.e.m., ^*^*p*< 0.05

## 4. DISCUSSION

By measuring neural oscillations in children with and without dichotic listening deficits, we show that amblyaudia (AMB) is characterized by (1) a stark (rightward) interaural asymmetry in dichotic listening performance; (2) stronger phase-locked β/γ oscillations within the bilateral AC for rapid stimuli presented to the right ear; (3) ear asymmetries in AC responses that predict the larger than normal right ear advantage in dichotic listening; and (4) weaker functional connectivity from right to left AC, despite overall stronger neural responses. Our findings reveal abnormally large auditory responses in children with AMB, particularly for right ear stimulation, which might lead functionally to a poorer behavioral capacity to integrate information between the two ears.

As far as we know, this is one of the first studies to conduct EEG TFA in children with dichotic listening deficits. Our data reveal unexpectedly larger β/γ AC oscillations (∼33 Hz) in children with AMB relative to their age-matched peers. Larger right ear responses across the board (Fig. 4) points to a pattern of ear-specific activation differences in AMB rather than one related to the cortical hemispheres, *per se*, corroborating previous fMRI work (Moncrieff et al., 2008). Previous neuroimaging studies have linked β-band activity with several perceptual-cognitive processes including working memory (Shahin et al., 2009; Zarahn et al., 2007) and auditory template matching (Bidelman, 2015, 2017; Shahin et al., 2009; Yellamsetty and Bidelman, 2018). Our task used only non-speech (repetitive click) stimuli and passive listening. Consequently, it is unlikely that the observed group differences reflect higher-level constructs associated with β frequencies (i.e., memory-based functions). Instead, our results better align with predictive coding/novelty detection (Cope et al., 2017; HajiHosseini et al., 2012; Sedley et al., 2016), sensory integration (Brovelli et al., 2004; von Stein and Sarnthein, 2000; Wang et al., 2017), and/or rhythmic prediction (Cirelli et al., 2014; Fujioka et al., 2009) accounts of β band oscillations. Under these latter interpretations, the higher β activity we find in the AMB group might be attributable to a relative inefficiency to integrate sensory information in the AC— perhaps more so if the repetitive stimuli used here (i.e., isochronous clicks) are treated by AMB listeners as quasi novel events.

Alternatively, β rhythms have been linked to inhibitory processing (Kropotov, 2010) which is often impaired in neurodevelopmental disorders (Milner et al., 2018). Thus, insomuch as β (and γ) auditory cortical oscillations reflect the brain’s ability to synchronize to external sounds (Baltus and Herrmann, 2016; Cirelli et al., 2014; Fujioka et al., 2009), the much larger ITPL responses in children with AMB may reflect a form of overexaggerated acoustic entrainment from reduced inhibitory processing that might arise from both bottom-up and top-down auditory pathways. Over-synchronization might diminish the ability to suppress irrelevant acoustic features, rendering less flexibility in how the brain adapts to changes in the sound environment. Such a mechanism based on aberrant (overly strong) auditory neural entrainment might account for at least some of the behavioral listening deficits observed in children with AMB.

Alternatively, if we consider the spectral differences to reflect canonical “γ activity,” this might reflect a different mechanism of perceptual-cognitive dysfunction between AMB and WNL listeners (Giraud and Poeppel, 2012). Induced γ activity has been attributed to high-level cognitive processes (Pulvermüller et al., 1997), the formation of perceptual objects, representations in auditory-lexical memory (Shahin et al., 2009), and integration of top-down and bottom-up auditory processing (Tallon-Baudry and Bertrand, 1999; Trainor et al., 2009). Enhanced γ in this study corroborates previous work suggesting γ activity might represent perceptual confusion, for example, when sounds do not conform to a unified identity (Bidelman, 2015). This might prevent higher perceptual learning that links neural representations to a perceptual output (Shahin et al., 2008). Early feature selection is also reflected by γ activity which can be modulated by attention (Gilley and Sharma, 2010; Mesgarani and Chang, 2012). γ activity is involved in the formation of coherent, unified percepts as neurons synchronize neural discharges across different brain regions (Buzsaki, 2006; Singer and Gray, 1995). In a study conducted in infants, left hemisphere γ reduction was seen for discrimination of segmental sublexical information at the phonemic level which contributed to information processing difficulties (Cantiani et al., 2019). However, in our study we find a right ear γ enhancement. It is thought that γ activity provides a building block of sensory-perceptual representations, and is hypothesized to have an indirect correlation with language outcomes (Cantiani et al., 2019; Yordanova et al., 2001). Thus, the increased γ responses we find in AMB may reflect a more rigid encoding of auditory perceptual objects which conceivably, may impair the ability to juggle complex sounds as required in dichotic listening and figure ground perceptual tasks. In children, γ band responses also reflect attentional modulation during sensory processing (Yordanova et al., 2001). Thus, another interpretation of increased γ activity in children with AMB could be due to a covert allocation of attention during auditory processing. Unnecessarily deploying attentional resources would make it more difficult to tune out (or tune into) information arriving at each ear. Still, we find this point speculative given the passive nature of our task.

Our results demonstrate that neural oscillations from the auditory cortex are unusually large in children with AMB, especially for right ear stimulation. This stark ear asymmetry in AC responses was predictive of behavioral scores; larger neural oscillations elicited from the right ear were correlated with larger asymmetries in dichotic word perception (as in children with AMB). Though neuroimaging data on APD is limited, these findings converge with recent EEG studies examining children with learning problems and auditory processing disorders. For example, Gilley et al. (2016) showed that in response to the speech, children with APD showed frequency and latency shifts specific to the upper-β/lower-γ frequencies, the same EEG bands identified here. They attributed these β/γ changes as reflecting a decoupling of early auditory encoding from the broader neural networks that govern auditory processing (Gilley et al., 2016). Cantiani et at. (2019) investigated neural substrates and oscillatory patterns of rapid auditory processing in infants with a familial risk of language and learning impairment and showed right-lateralized responses in θ and γ bands suggesting hemispheric differences in underlying oscillatory activities (Cantiani et al., 2019). Also, Yordanov et al (2001) conducted research to compare γ band in ADHD children with their normal peers to investigate attention-related differences between groups and reported ADHD children had larger amplitude and stronger phase-locked γ oscillations to right ear stimuli. They suggested an alteration of early auditory processing as a result of impaired motor inhibition (Yordanova et al., 2001). Collectively these studies show aberrant, over-arousing responses and/or altered pattern of habituation to stimuli affects mainly high-frequency oscillations (β and γ) that might be associated with bottom-up processing (Buzsáki et al., 2013; Friederici and Singer, 2015). However, since β and γ bands have been attributed to top-down and attentional processing, it is hard to fully rule out top-down explanations.

Several ERP studies have investigated the neural basis of dichotic listening. The time course, location, and extent of brain activation differ according to various dimensions of dichotic listening tasks (Jerger and Martin, 2004). For example, using binaural paradigms, Jerger et al (2004) showed poorer ERPs in APD children over left brain regions for speech stimuli but poorer responses over right regions for non-speech stimuli (Jerger et al., 2004; Jerger et al., 2002). In our own ERP work (Moncrieff et al., 2004), children with dichotic left ear deficits showed increased latencies, decrease amplitudes, and lateralized scalp distributions, implying slower neural conduction times, decreased interhemispheric transfer, and left ear failure to suppress competing signals arriving at the right ear, all of which could be indicators of binaural integration deficits (Moncrieff, 2006). However, such scalp effects are not always easily interpretable. Electrode waveforms, for instance, may reflect the involvement of frontal generators known to contribute to the auditory ERPs (Bidelman and Howell, 2016; Knight et al., 1989; Picton et al., 1999), especially during dichotic listening tasks (Bayazıt et al., 2009). Our use of passively presented, monaural stimuli helps rule out the confounds of linguistic stimuli and attention which modulate hemispheric and ear asymmetries. Moreover, the use of source analysis provides a novel window into the nature of DL deficits by identifying neurophysiological changes in AMB listeners localized to early auditory cortical areas.

The present time-frequency and connectivity data corroborate but extend these previous findings. Functional connectivity revealed that while LH→RH transmission was identical between groups, children with AMB showed weaker transfer of information from right to left AC, despite their overall stronger neural responses. These findings confirm a previously posited interhemispheric transmission deficit in AMB (e.g., Moncrieff et al., 2008, p. 43), routed toward the linguistic (left) cerebral hemisphere. Presumably, the weaker right-to-left transfer we find in AMB patients is grounded in interhemispheric connections via the corpus callosum (CC). Indeed, Clarke et al (1993) found that RE scores were negatively correlated with anatomical CC size; patients with larger interhemispheric pathways did more poorly in RE performance. The authors used dominant and non-dominant scores to remove the effects of those with better performance in their LE and posited that better RE performance stemmed from a release from inhibition following split-brain surgery; the RH could no longer interfere with LH processing (Clarke et al., 1993). In the current study, the monaural nature of our results suggests there is a failure in suppression which could occur either subcortically or within the CC on the side of the RH. Our connectivity findings also link to the “structural theory” of DL (Kimura, 1967) where the ipsilateral pathway is assumed to play a suppressive role and is weaker in AMB. However, the degree to which the ipsilateral pathway is inhibited (and therefore contralateral dominates) is unknown in our (nonspeech) monaural data but could be tested in future experiments with dichotic (binaural) tasks (cf. Della Penna et al., 2007).

Using EEG connectivity, we have previously shown that β band transmission to the AC increases with other forms of hearing deficits (i.e., age-related hearing loss; Price et al., 2019). Thus, the fact that β-γ connectivity also differed in children with AMB in the present study suggests these EEG frequencies may indeed reflect a decoupling of auditory sensory processing from the brain networks that subserve perception (Gilley et al., 2016). That these functional changes have not observed in lower EEG frequencies (e.g., θ or α band; < 10 Hz) (Gilley et al., 2016) further implies the neural effects in AMB listeners are not due to broad, domain-general cognitive mechanisms (e.g., attention). This is further bolstered by the fact we observed group differences *within* the early AC, for *passive* listening, and *nonspeech* stimuli. Taken together, emerging EEG evidence (present study; Cantiani et al., 2019; Gilley et al., 2016) argues against a high-level, linguistic, or attention-based account of AMB/APD.

A right ear advantage can be seen both in normal and APD children, particularly in the form of poorer verbal processing of dichotic sounds (present study; Bellis, 2011; Bryden, 1963; Kimura, 1961; Satz et al., 1965). It has been hypothesized that cognitive factors such as attention and short-term memory may influence the strong bilateral asymmetries in RDDT performance among APD children (Moncrieff and Wilson, 2009). While these factors might impact behavioral ear asymmetries, they cannot account for neural differences nor the brain-behavior link between the click-evoked oscillations and dichotic listening score laterality seen here (Fig. 5). Our results lead us to infer anomalous asymmetries in children with AMB are due instead to a lack of appropriate sensory processing (e.g., reduced inhibition, overexaggerated neural entrainment, and/or poorer cross-talk between auditory cortices). We hypothesize that when sound is presented to the right ear, listeners with AMB fail to inhibit (or over-represent) those signals due to a deficiency of inhibitory interneurons in both hemispheres or the auditory brainstem (Pfurtscheller et al., 1997), resulting in abnormal asymmetry/laterality between the ears. Still, why brain-behavior associations between oscillations and dichotic listening are restricted to the DWT test is unclear. In contrast to DWT, RDDT and CW might place higher demands on working memory and attention, respectively (Moncrieff, 2011). This could be why we failed to observe correlations between these two tests and brain responses. At the very least, this speaks to the critical need to consider multiple dichotic listening assessments for amblyaudia evaluation (Moncrieff et al., 2016). It is also conceivable that neural asymmetries in listeners with AMB originate prior to AC. Functional ear asymmetries have been reported in brainstem frequency-following responses (Ballachanda et al., 1994; Krishnan et al., 2011), which are predictive of asymmetrical processing in later cerebral cortex (e.g., Efron, 1985). Future studies are needed to evaluate potential brainstem correlates of the clinical manifestations of AMB.

## Acknowledgments

We thank Drs. Karen Bell and Caitlin Price for comments on earlier versions of this manuscript. Requests for data and materials should be directed to G.M.B. This work was supported by the National Institute on Deafness and Other Communication Disorders of the NIH under award number R01DC016267 (G.M.B.), the U.S. Department of Education under award number H325K100325 (D.M.), and the Lions Hearing Research Foundation (D.M.).

## Footnotes

1 The online high pass filter (>10 Hz) during data acquisition meant there was negligible induced activity in our data. Induced activity is also difficult to measure in paradigms with rapid stimulus presentation as used here (Price et al, 2019). Thus, we decided to consider only phase-locked oscillations (i.e., ITPL) in our analyses.

